# Visualizing the cytosolic delivery of bioconjugated QDs into T cell lymphocytes

**DOI:** 10.1101/2020.09.12.294991

**Authors:** Haoran Jing, Marcell Pálmai, Badeia Saed, Anne George, Preston T. Snee, Ying S. Hu

## Abstract

The aggregation state and endosomal trapping of engineered nanocarriers once internalized into cells remain poorly characterized. Here, we visualized the membrane penetrating dynamics of semiconductor quantum dots (QDs) into the cytosol of T cells on a single-cell and single-nanoparticle basis. We water solubilized CdSe/CdZnS QDs with polymer encapsulants functionalized with a cell-penetrating peptide composed of an Asp-Ser-Ser (DSS) repeat sequence. T cells tolerated the 24-h incubation with QDs at concentrations of 5 nM or lower. Single-particle imaging demonstrated that the number of internalized nanoparticles was dependent upon the concentration of the probes for both control (peptide-free) and DSS-QDs. DSS-QDs were mostly distributed as monomers, whereas the control QDs were aggregated into clusters. Single-particle tracking using total internal reflection and highly inclined illumination showed that DSS-QDs were stationary near the activating surface and mobile within the cytosol of the T cell. A correlation exhibited between the mobility and aggregation state of individual QD clusters, with monomeric DSS-QDs showing the highest mobility. In addition, monomeric DSS-QDs displayed much faster diffusion than the endosomes. A small-molecule endosome marker confirmed the absence of colocalization between endosomes and DSS-QDs, indicating their endosomal escape. The ability to deliver and track individual QDs in the cytosol of live T cells creates inroads for the optimization of drug delivery and gene therapy through the use of nanoparticles.

## Introduction

Biotechnology and biomedical applications of nanomaterials have flourished in the last decade, especially in the fields of biomaterial separation, immunoassays, diagnostics, and drug delivery systems (1-6). To enhance their diagnostic and therapeutic efficacy, novel nanoassemblies must be engineered to function in biologically relevant environments and have multivalent loading capacity to facilitate detection and effective drug delivery. Reproducibly accessing the intracellular space with precision delivery has remained a highly desirable goal. In particular, harnessing the cellular machinery to suppress (7) or enhance (8) the cellular function for treatment using theranostic nanoparticles is a highly promising strategy.

T cell lymphocytes have been engineered as “programmable and living drugs” for the treatment of various types of refractory cancers (9). To this end, engineering nanoparticle carriers for drug and gene delivery to targeted T cell populations represents a promising strategy for increasing the response rate and durability of immunotherapies. The unique physical, photophysical, and photochemical properties of engineered nanomaterials have enabled a broad range of finely controlled stimulation of T cells. For instance, magnetic nanoparticles have been used to enhance T cell activation *via* forced spatial clustering of T cell receptors and co-stimulatory molecules using an external magnetic force (10). Photothermal and photodynamic therapy using nanoparticles have shown efficacy towards augmenting the anti-tumor response and T cell infiltration (11-13). Upconverting nanoparticles enable the delivery of visible light into deep tissues and remote-controlled immunomodulation using ultraviolet light-activatable immunostimulatory agents based on CpG oligonucleotides (14). In addition, targeted delivery of small molecule inhibitors to specific T cell subpopulations effectively reduces the toxicity of systematic administration and improves antitumor immunity (15).

Transmembrane receptors with stimulatory or inhibitory functions, such as the mechanisms targeted by the immune checkpoint inhibitors, represent a major class of targets for engineered nanoparticles. A complementary approach involves modulating the intracellular signaling pathways by delivering small molecule drugs (10, 16) or genetic materials such as siRNAs (17-20). This requires the delivery of engineered nanoparticles into the intracellular space. However, the delivery and localization of nanoparticles within cells have been mainly characterized using transmission electron micrographs (TEM) of fixed samples (10) or confocal imaging of particle ensembles (21). Compared to fluorescence imaging, TEM has limited capability in multiplexed labeling and the detection of biomolecules. Moreover, systematic investigations of how design parameters affect the nanoparticle-T-cell interactions and subsequent cell entry require high-resolution live-imaging assays. However, confocal imaging lacks the necessary spatial resolution, and single-molecule imaging is difficult to perform with suspension T cells.

Semiconductor quantum dots (QDs) have excellent characteristics, such as size-tunable optical properties and photochemical stability, for single-particle visualization in live T cells (22). QDs exhibit high molar extinction coefficients over a broad excitation region and narrow emission spectra, enabling multiplexed detection from the blue to near-infrared region (23). The development of methods for robust and reproducible cytosolic delivery of QDs into live cells has been challenging for a variety of reasons (24). Generally, the highest quality nanomaterials are prepared in a hydrophobic solvent and require surface modification to impart water solubility. Surface functionalities are often composed of amines, carboxylic acids, or PEG grafted onto polymers, liposomes or small molecule “caps” to name a few (25-29). Unfortunately, without further functionalization, these nanomaterials have minimal interactions with cells or may at best become trapped inside of endosomes (30, 31). To date, the majority of reports on successful cytosolic delivery use protein functionalized QDs to either circumvent endosomal trapping or escape once sequestered (31-34). Other strategies involve mechanical delivery, such as microinjection (7), which may be difficult to implement on T cells. Recently, cell-penetrating poly(disulfide)s have been reported to be effective in drosophila cells (35). However, the number of reports on cytosolic delivery of fluorescent QDs remains few and far between, and it remains unclear what surface modification parameters can be optimized for maximal cargo delivery, and how such nanoparticles may interact with human T cells.

Here, the cytosolic delivery of biocompatible QDs into T cells was investigated. Cell uptake of CdSe/CdZnS QDs was enhanced by surface coating with a cell penetrating peptide (CPP) composed of a repeated Asp-Ser-Ser (DSS) sequence. The DSS repeats are derived from a motif found in dentin phosphophoryn, one of nature’s most acidic proteins, which was previously demonstrated by us to deliver quantum dots into live cells (21). Furthermore, the DSS motif was used to coat lignin nanoparticles, facilitating their efficacy as a drug delivery agent for several types of cancer cells (36). To enable stable single-particle imaging, QD-laden T cells were immobilized on activating surfaces. Control and DSS-QDs were found to be distinctively distributed between the monomeric and clustered state, with the majority of DSS-QDs in the single-particle state. In addition, DSS-QDs displayed significantly higher mobilities compared to control QDs. This study creates in-roads for the use of DSS-coated nanoparticles for *in vivo* drug delivery and gene therapy.

## Results

Our previous study demonstrated that dentin phosphophoryn (DPP), an acidic, phosphorylated protein that is a ubiquitous component of the dentin extracellular matrix, is internalized by several cell types *via* a non-conventional endocytic process (21, 37). Furthermore, as DPP contains Asp-Ser-Ser (DSS)_n_ repeats distributed throughout the protein, it was demonstrated that (DSS)_n_ facilitates endocytosis and can function as a cell-penetrating peptide to deliver proteins for therapeutic applications (21, 36). DSS can internalize cargo such as CdSe/CdZnS quantum dots in several cell types. QDs conjugated with the chimeric protein DSS and the osteoblast-specific transcription factor Runx2 (DSS-Runx2) favored nuclear translocation (21). Incubation of the functionalized nanoparticles with MC3T3 osteoblast precursor cells resulted in passive delivery into the cytoplasm and trafficking into the nucleus. As DPP contains Asp-Ser-Ser (DSS)_n_ repeats distributed throughout the protein, it was demonstrated that (DSS)_n_ facilitates endocytosis and can function as a cell-penetrating peptide to deliver proteins for therapeutic applications (21, 36). In the present study, a linkable (DSS)_10_K_4_ polypeptide was synthesized and conjugated to water-soluble CdSe/CdZnS QDs to evaluate their membrane penetration activities with T cells. E6-1 Jurkat T cells were incubated with the growth medium containing DSS-QD solution for 24 h. Before imaging, T cells were washed three times in HBSS and resuspended in phenol red-free HBSS containing 1% FCS. The T cells were immobilized on an 8-well chamber slide coated with anti-CD3 antibodies (clone OKT3). QD-laden T cells were subsequently imaged using TIRF with a Nikon Eclipse Ti-E2 inverted microscope (**Fig. 1**) (38).

**Fig. 1.**
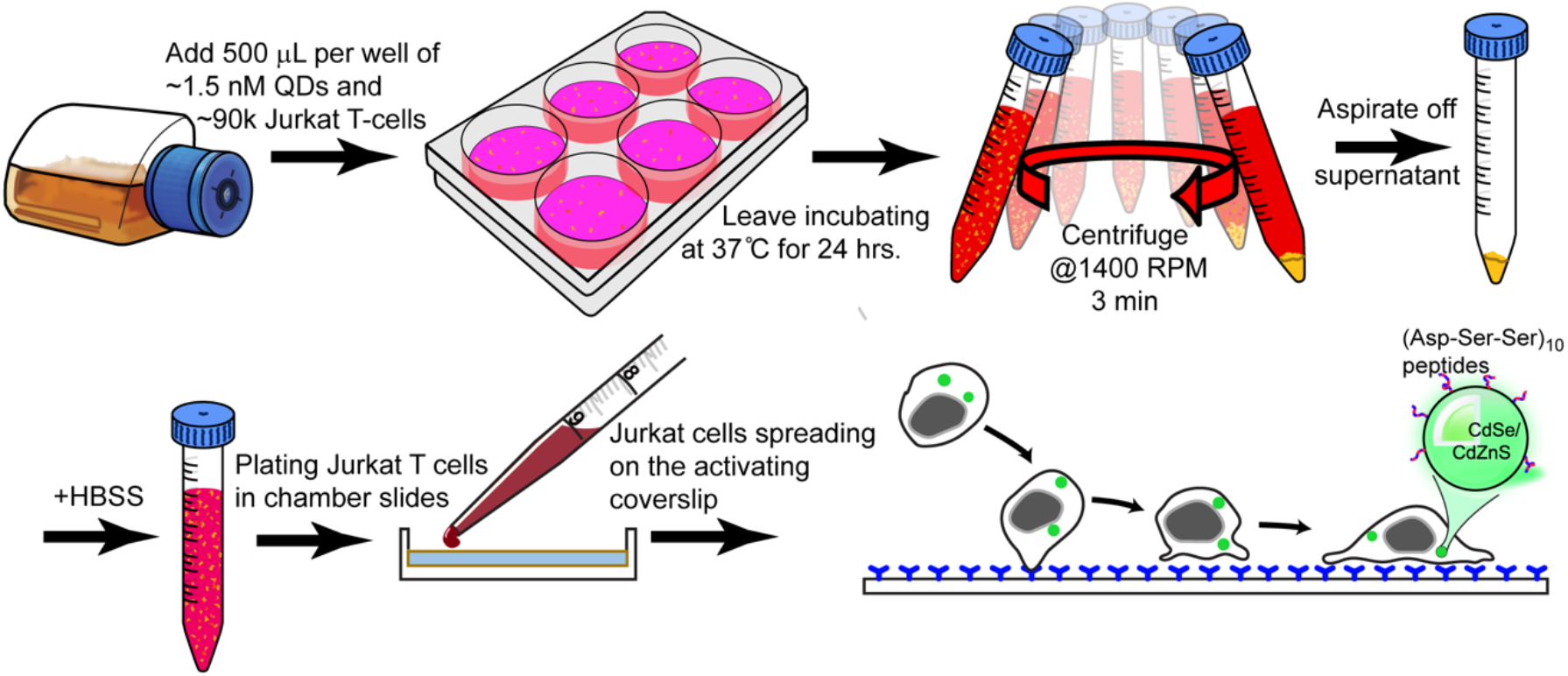
Schematic representation of the T cell incubation with QDs and subsequent washing, immobilization, and imaging on the activating surface coated with antibodies.

To evaluate whether the internalization of DSS-QDs affected T cell activation, both the immobilization and activation process, Jurkat T cells were examined by brightfield and fluorescence imaging. **Fig. 2*A*** shows a QD-laden T cell initially interacted with the OKT3-coated surface and then maintained a stable contact with surface. On the activating surface, brightfield imaging revealed no morphological differences between DSS-QD-laden T cells and control T cells without prior QD exposure. Notably, DSS-QDs were visible inside the T cells through TIRF imaging (**Fig. 2*B***). To evaluate the cytotoxicity of the QDs terminated with DSS peptide on Jurkat T cells, a CellTiter-Glo^®^ Luminescent Cell Viability Assay was conducted on cells with DSS-QDs for 24h. **Fig. 2*C*** shows that the cell viability remained above 80% over the applied QD concentration range from 3.16 pM to 10 nM. The IC50 of DSS-QD was found to be 23.2 nM. These results reveal that DSS-QDs have no considerable cytotoxicity under this concentration range and support the suitability of DSS-QDs for studying nanoparticle internalization into T cells.

**Fig. 2.**
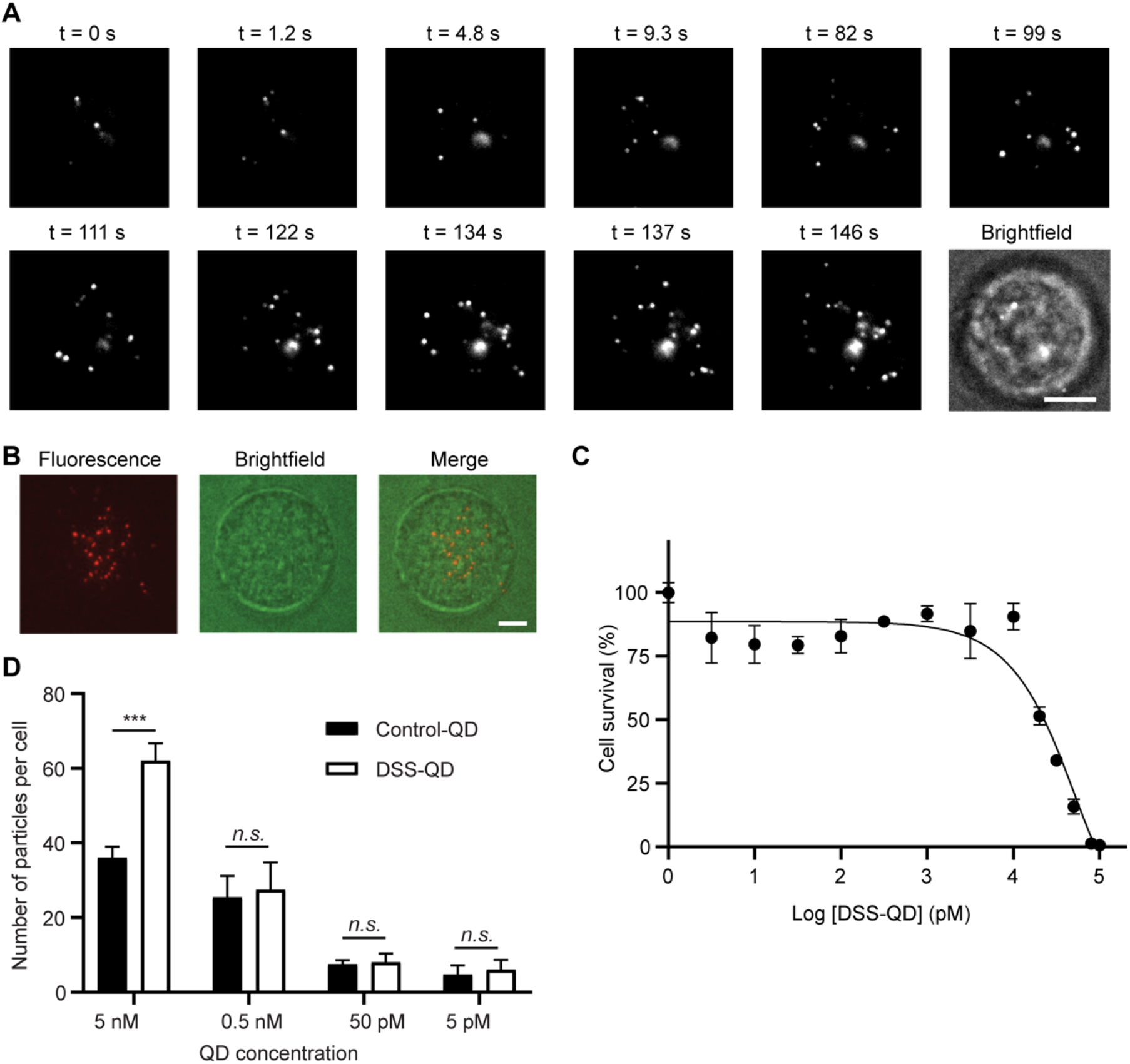
Single-particle visualization of QD delivery into T cell lymphocytes. (A) Selected imaging frames showing the initial interaction between a T cell and the activating surface coated with anti-CD3 antibodies. (B) Brightfield and fluorescence images of a Jurkat T cell on the activating surface demonstrating the QD delivery into the cell. (C) Representative IC50 results for incubating Jurkat T cell for 24h with DSS-QDs in the culture medium. Each data point represents the mean ± standard deviation of wells performed in quadruplicate. (D) Quantification of the number of internalized DSS-QDs per cell characterized by TIRF imaging. Statistical significance was evaluated using an unpaired Student’s t-test. *** represents *p<*0.001 and *n*.*s*. denotes not significant. Scale bars: 5 µm.

A brief examination of the control and DSS-QD images revealed that the internalization of nanoparticles was concentration-dependent. An ImageJ plugin, ThunderSTORM, with multiple-emitter-fitting capability was used to quantify this observation (39, 40). **Fig. 2*D*** shows that the number of observed QDs per cell monotonically decreased when the concentration of the QDs was reduced. Between the two groups, DSS-QDs showed significantly more internalization when the T cells were incubated with a high concentration of 5 nM DSS-QDs in the growth medium. On average, the number of internalized QDs per cell was 62 ± 9 (mean ± SEM, *n* = 36 cells) for DSS-QDs and 38 ± 6 (mean ± SEM, *n* = 28 cells) for control QDs. This difference was diminished as the concentration decreased (**Fig. 2*B***). Furthermore, control and DSS-QDs revealed distinct particle distributions in the form of large and small clusters.

**Fig. 3*A* and *B*** are typical images of activated T cells after 24 h incubation with DSS-QDs and the control materials. At the 0.5 nM concentration, the random distribution of DSS-QDs is manifested as sparse and small dots inside the cell, while the control-QDs displayed a more localized distribution and pronounced puncta. Incubation with a higher QD concentration of 5 nM increased the number of internalized DSS-QDs, while pronounced clusters of control QD were still visible. To investigate whether the variation in the brightness resulted from different optical behaviors of control *vs*. DSS-QDs, single-particle imaging was performed on a coverglass coated with monodispersed QDs. **Fig. 3*C*** demonstrates a background-corrected average brightness value of approximately 40-60 a.u. for both QDs at the single-particle level). **Fig. 3*D*** plots the intensity profile of internalized QD clusters. The majority of DSS-QDs registered intensity levels below 80 a.u., indicating that DSS-QDs were mostly monomeric in the T cell cytosol. In contrast, the intensity profile of control QDs was much more heterogeneous; the intensity distribution displayed a minor peak between 160 and 480 a.u. and a major peak greater than 1040 a.u. Based on the single-particle intensity, the number of QDs ranges from 4 to 12 in small aggregates and exceeds 20 in large aggregates.

**Fig. 3.**
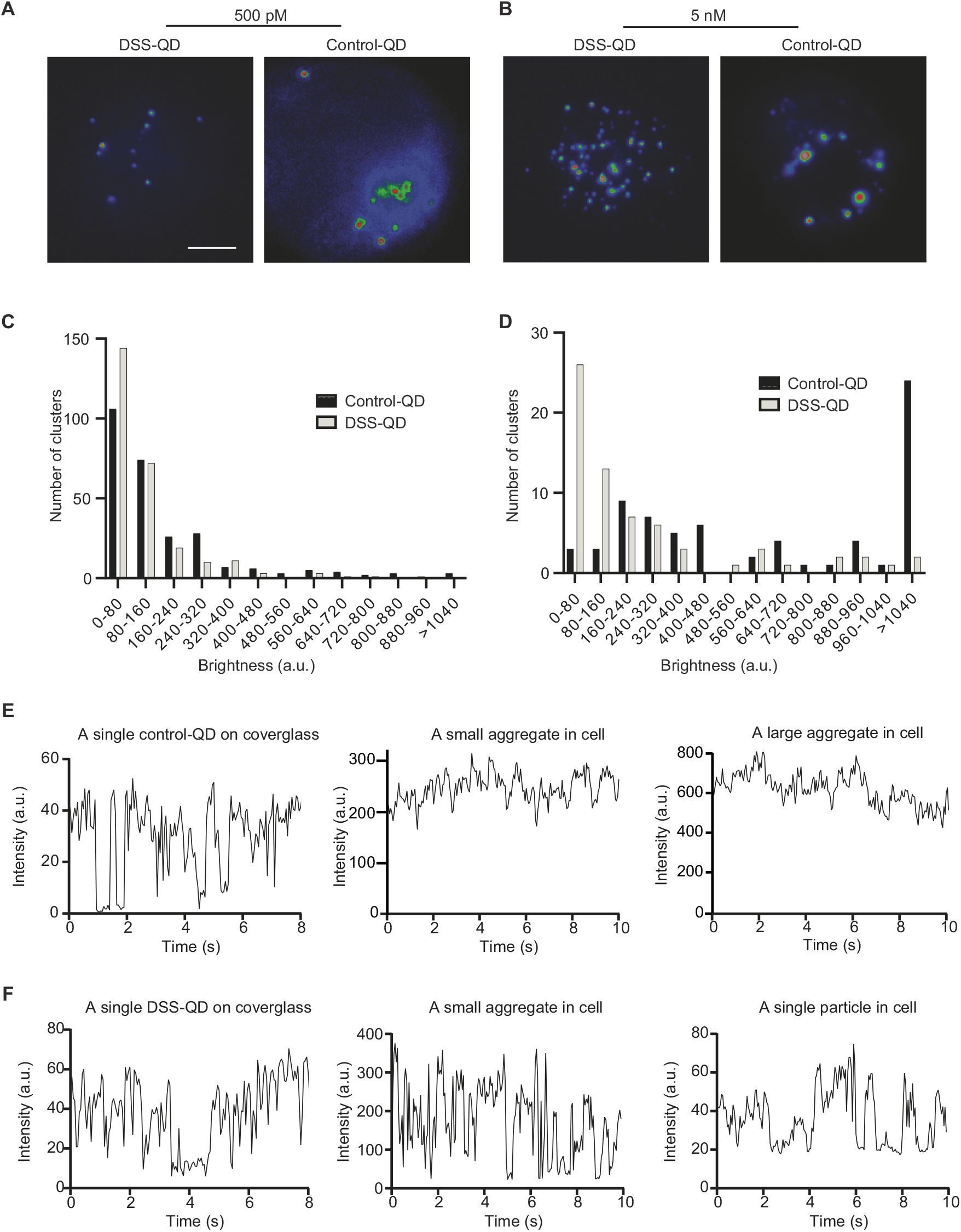
Intensity analyses indicate distinct aggregation states of DSS-QDs and control-QDs in the cytosol of T cells. A) Fluorescent images of internalized DSS-QDs and control-QDs after 24 h incubation with 0.5 nM of the QDs. Scale bar: 5 µm. B) Fluorescent images of internalized DSS-QDs and control-QDs after 24 h incubation with 5 nM of the QDs. C) Intensity profiles of individual QDs dispersed on a coverglass. D) Intensity profiles of QD clusters inside T cells. E) Time traces of emission intensities of control-QDs on the coverglass and in the T cell cytosol. F) Time traces of emission intensities of DSS-QDs on the coverglass and in the T cell cytosol. Images were acquired under identical experimental conditions.

Due to the fluorescence intermittency of CdSe QDs, a random switching takes place between bright fluorescence periods (“on” state) and dark, non-emissive ones (“off” state). The sharp transition between the “on” and “off” state is visible from time trace plot of single QDs on the cover glass (left panels in **Fig. 3*E* and *F***). Clusters of QDs exhibited fluctuations of intensities at much higher levels. Small dots of DSS-QDs in the T cell cytosol displayed identical blinking properties compared to single QDs on the cover glass (**Fig. 3*F***), further confirming their monomeric state.

To investigate the dynamics of internalized QDs and how particle dynamics correlate with the cluster size, two-dimensional single-particle tracking was performed. Intensity profiling was utilized to categorize the aggregation state of each cluster into a single particle (approximately 40-60 a.u.), a small aggregate (< 500 a.u.), or a large aggregate (> 500 a.u.). A similar approach has been previously used to quantify membrane receptors *via* calibrated QD intensities (41). **Fig. 4*A*** shows that changes of single-particle positions can be detected between image frames 50-ms apart. These observations prompted a changeover in illumination to a highly inclined and laminated optical sheet (HILO) configuration. **Fig. 4*A*** illustrates representative single-particle images and tracks of control and DSS-QDs after 24-h incubation with 5 nM QDs. For control-QDs, the large aggregates displayed significantly more confined diffusion than the small aggregates (red *vs*. black track). Most DSS-QDs were dispersed in the cytosol as monomers with the emergence of small aggregates at higher QD concentrations. Monomeric QDs were mobile in the cytosol (green track). Averaged two-dimensional mean squared displacement (MSD) showed similar characteristics (**Fig. 4*B***), The diffusion coefficient of single DSS-QDs was found to be 7.5 ± 1.9×10^−2^ μm^2^/s. The linear MSD curves indicate normal diffusion observed within the imaging duration of approximately two seconds.

**Fig. 4.**
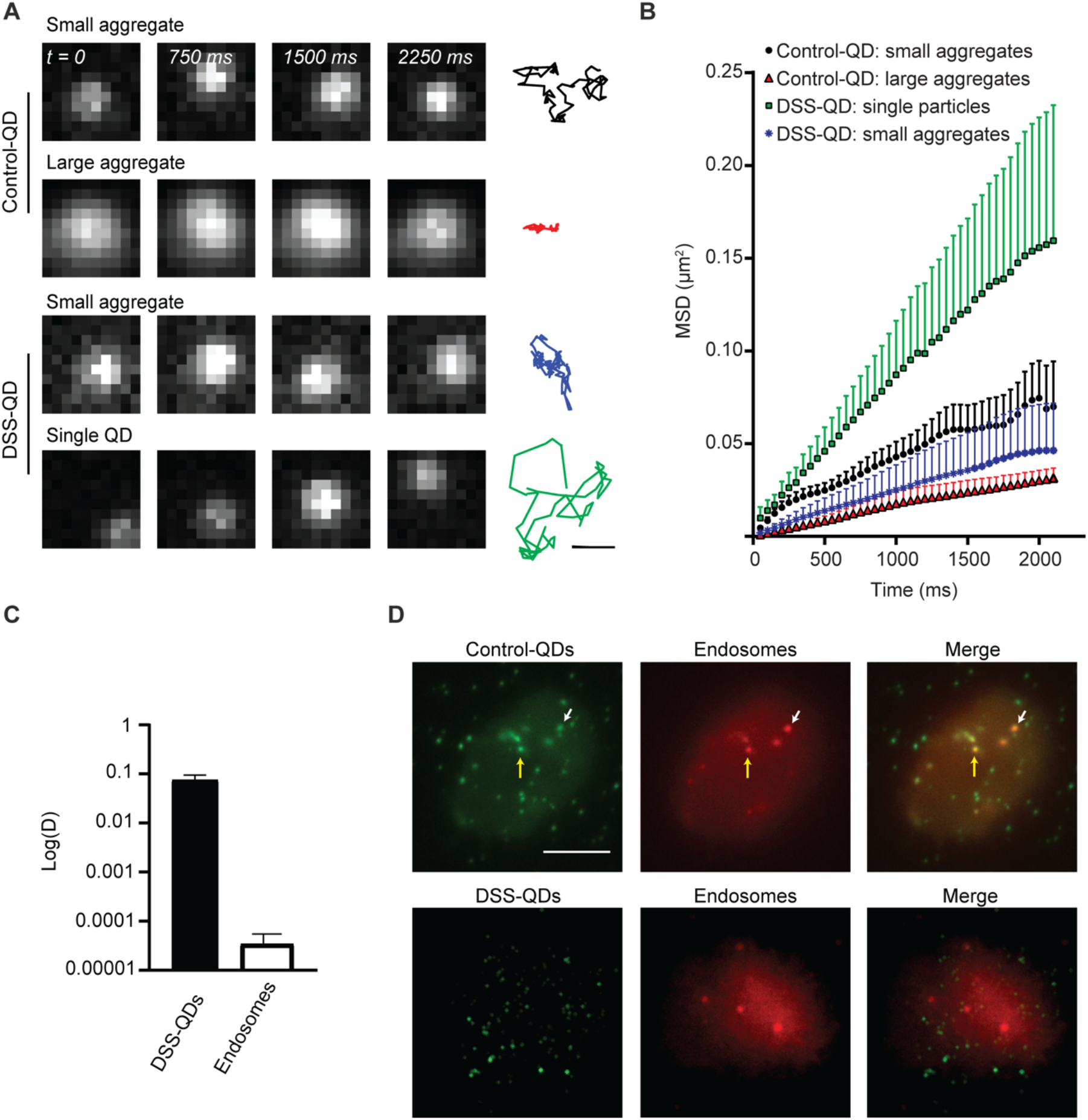
Two-dimensional single-particle tracking and fluorescence imaging using an endosomal marker reveal that DSS-QDs were not entrapped in endosomes. A) Single-particle dynamics of QDs imaged by TIRF. Left panel: Single-particle imaging, Right panel: Representative single-particle tracks of small and large aggregates of control QDs (black and red, respectively), and small aggregates and single particles of DSS-QDs (blue and green, respectively). Scale bar: 300 nm. B) Averaged MSD measurements with the error bars indicating the standard error of mean. Error bars are shown only above the mean for clarity. C) Diffusion coefficient log_10_ value of the diffusion coefficient D in µm^2^/s for single DSS-QDs and endosomes. D) Fluorescence images of an T cell labelled with QDs and acridine orange demonstrating the control-QDs delivery into endosomes. White and yellow arrows mark the colocalization of control-QDs and endosomes. Scale bar: 5 µm.

Next, T cells were incubated with the small-molecule dye acridine orange (AO) to label acidic endosomes. Through single-particle imaging, the diffusion coefficient of endosomes was found to be 3.7 ± 2.2×10^−5^ μm^2^/s. These data are consistent with **Fig. 4*C*** that demonstrates slower endosomal diffusion compared to monomeric DSS-QDs in T cells. Studies by others reported a higher diffusion coefficient for actively trafficking endosomes. These values range from 2.1 ± 0.1×10^−3^ μm^2^/s for kinesin-directed endosome diffusion during cell division (42) to 2.4 ± 0.7×10^−2^ μm^2^/s in the motile population during early endosome trafficking (43). Of note is that the diffusion coefficient of DSS-QD is still higher than these values. The rapid diffusion suggests that DSS-QDs are free from endosomes whereas large aggregates of control QDs are likely sequestered within. We further confirmed these results by co-incubating T cells with QDs and the AO dye. **Fig. 4*D*** shows that, while several pronounced puncta of control QDs overlapped with the endosome vesicles (arrows in **Fig. 4*D***), no obvious overlap between the DSS-QDs and endosomes was observed.

## Discussion

Our results establish a methodology to evaluate cell penetrating behaviors of individual nanocarriers using high resolution microscopy. We observed internalization of both control and DSS-QDs into T cells from the culture medium. The observation is aligned with the notion that T cell lymphocytes may exhibit some basal levels of phagocytic activities. For instance, human γδ T cells have been reported to be capable of professional phagocytosis (44). T cells may also internalize QDs through micropinocytosis facilitated by the microvilli structures on the plasma membrane (45). Importantly, our data provide the evidence of QD internalization in the low-concentration range, which may be below the detection threshold of standard imaging techniques, such as confocal microscopy.

In the present study, we observed different characteristics between control and DSS-QDs inside T cells (**Fig. 3 and 4**). The brightness and photoblinking behaviors of QDs confirm the presence of monomeric DSS-QDs inside the cell. Through single-particle tracking, DSS-QDs were observed to diffuse at much faster speeds than that of endosomes. Control QDs were found to be in an aggregated state and likely trapped within endosomes, which suggests that they experience a different cell entry mechanism. We anticipate that single-particle imaging can be applied to further investigate these mechanisms across different cell types.

Our imaging study is limited to regions near the basal membrane of the T cell. As QD-laden T cell started to interact with the activating surface, intracellular QDs emerged in the focal plane and remained highly dynamic (**Fig. 2*A***). To fully characterize QDs in the intracellular space, single-particle volumetric imaging, such using the lattice light-sheet system (46, 47), may be utilized. In addition to the particle concentration, the temperature, pH, and incubation duration may also affect the QD characteristics in the cytosol and are subjects of future investigations.

In summary, we investigated the cytosolic delivery of DSS-QDs in suspension T cells and the aggregation state and dynamics of internalized QDs on a single-particle basis. Functionalized CdSe/CdZnS QDs were synthesized and water solubilized with amphiphilic poly(acrylic acid) functionalized with a peptide cell delivery vector comprised of a repeat sequence of aspartic acid and serine. A protocol for immobilizing T cells on activating surfaces was developed to enable stable single-particle tracking of QDs. DSS-QDs were found to be monomeric or in small clusters. In comparison, control QDs without the DSS peptide were generally aggregated. Through single-particle tracking, monomeric DSS-QDs showed substantially higher mobility compared to the small-cluster counterparts and the larger clusters of control QDs. Our imaging data also revealed the colocalization of endosomes and control-QD clusters in the T cells and absence of colocalization for DSS-QDs. The imaging platform enables systematic investigations of design parameters that affect nanoparticle-T-cell interactions at the level of single particles in vivo, thereby advancing the understanding of nanomaterials dynamic in T cell cytoplasm and their use for enhancing cancer immunotherapy.

## Materials and Methods

### Materials

Gibco RPMI 1640 (Cat# 11875093), Gibco fetal bovine serum (Cat # 10437036), Gibco Hank’s Balanced Salt Solution (HBSS) (Cat# 14-025-092), Lab-Tek™ 8-well chambered coverglass (Cat# 155409), Acridine Orange, 10mg/mL in water, (Invitrogen, Cat # A3568), and Nunc™ 12-well plate (Cat# 150628) were purchased from Thermo Fisher Scientific (USA). Fetal bovine serum (Cat# F0926), bovine serum albumin (Cat# 5470-1G), CaCl_2_, MgCl_2_ were purchased from Sigma-Aldrich (St. Louis, MO, USA). In vivo Mab OKT3 antibody (Cat# BE0001-2-25MG, Lo. 683618M2) was purchased from BioXCell (Lebanon, NH, USA). Poly(acrylic acid), average M_W_ 1800, was purchased from Aldrich. N-octylamine was from TCI and EDC was purchased from G-Biosciences. K_4_(DSS)_10_ was prepared by the UIC Research Resource Center and was purified by HPLC prior to use. Hydrophobic, tetradecylphosphonic acid coated CdSe/CdZnS QDs were prepared according to a previously published procedure (48).

### Control QD and DSS QD synthesis

Control QDs were water solubilized using 40% octylamine-modified poly(acrylic acid) as per the protocol outlined in ref. 47. To prepare DSS-QD conjugates, initially water soluble QDs and DSS peptide were incubated with poly(ethylene glycol) conjugation reagent (33, 49). However, it was found that the resulting materials did not display significantly different behavior compared to controls. The preparation was modified to enhance the yield of functionalized nanomaterials by conjugating DSS to the modified acrylic acid solubilizing polymer first, which was then processed by precipitation in acidic water and subsequently used to solubilize the CdSe/CdZnS nanomaterials as outlined below.

### DSS QD conjugation

Our approach is to conjugate DSS peptide to the modified acrylic acid solubilizing polymer first, which was then processed by precipitation in acidic water and then used to solubilize the CdSe/CdZnS nanomaterials. To this end, 5 mg of (DSS)_10_K_4_ (1.46 mmol) was added to a solution of DMF with 8 mg poly(acrylic acid) (111 mmol) and 9 mg EDC (47 mmol). After stirring a few moments, 7.3 μL of octylamine (44.1 mmol) was slowly added. The sample was stirred overnight and was precipitated with the addition of water. After centrifugation, the supernatant was discarded. The functionalized polymer was dissolved in basic water, and was titrated with mildly acidic water to <pH 5, upon which the polymer precipitated. The supernatant was discarded, and the polymer was dried under vacuum. The final mass was 12 mg (64% yield). Approximately 6 mg was used to solubilize 1.8×10^−8^ moles CdSe/CdZnS QDs as previously reported (50).

### QD incubation with Jurkat cells

Jurkat E6-1 cells were obtained from ATCC. For the incubation, 500 *μ*L of 5 µM QDs was added with ∼90 k Jurkat cells in culture medium (RPMI, 10% FCS) to a 12 wells plate. Jurkat cells were cultured with QD-containing medium at 37 °C and 5% CO_2_ for 24 hours. Jurkat cells were then transferred into a 10 mL conical tube with 5 mL HBSS added into the medium and centrifuged at @1400 rpm for 3 min for three times. The cells were resuspended in the imaging buffer containing 1% bovine serum albumin, 0.5 mM Ca^2+^, 2 mM Mg^2+^, and HBSS pre-warmed to 37 °C.

### Acridine Orange (AO) incubation with Jurkat cells

Before imaging, Jurkat cells were removed from a culture flask and placed in a 1.5 mL centrifuge tube. They were incubated in 6.6 nM Acridine Orange (AO) in culture medium for 15 min at 37 °C in the dark. The 1.5 mL centrifuge tubes were wrapped in aluminum foil to protect the dye from light exposure. After incubation, cells were washed with 1.5 mL Gibco Hank’s Balanced Salt Solution (HBSS) three times, and then were resuspended in DPBS.

### Immobilization and imaging of T cells

To make the activating surface, 8-well chamber slides were cleaned with absolute ethanol and DI H_2_O, then incubated overnight at room temperature. Coated OKT3 surface was produced by adding 200 μL OKT3 antibody at a concentration of 1 μg/mL in PBS per well. TIRF was performed on a Nikon N-Storm super resolution ECLIPSE Ti2-E microscope (TIRF 100 ×, 1.49 NA objective lens). QD-laden T cells were a 405 nm continuous wave laser and the emission was collected at 561 nm. The images were collected by a Photometrics Prime 95B sCMOS camera with a pixel size of 110 nm. Image analysis was performed by the ThunderSTORM ImageJ plug-in and a custom MATLAB code.

## Acknowledgments

The authors would like to thank the Department of Chemistry at the University of Illinois at Chicago for their support.

## Author Contributions

Y.H. and P.T.S. conceived the experiment, A.G., M.P. and P.T.S synthesized CdSe/CdZnS QDs and DSS-QDs, H.J. performed the T cell and imaging experiments, H.J. and Y.H analyzed the imaging data, all authors contributed to the writing and editing of the manuscript.

